# A double dissociation in sensitivity to verb and noun semantics across cortical networks

**DOI:** 10.1101/319640

**Authors:** Giulia V. Elli, Connor Lane, Marina Bedny

## Abstract

What is the neural organization of the mental lexicon? Previous research suggests that partially distinct cortical networks are active during verb and noun processing. Are these networks preferentially involved in representing the meanings of verbs as opposed to nouns? We used multivoxel pattern analysis (MVPA) to investigate whether brain regions that are more active during verb than noun processing are also more sensitive to distinctions among their preferred lexical class. Participants heard four types of verbs (light emission, sound emission, hand-related actions, mouth-related actions) and four types of nouns (birds, mammals, manmade places, natural places). As previously shown, the left posterior middle temporal gyrus (LMTG) and inferior frontal gyrus (LIFG) responded more to verbs, whereas areas in the inferior parietal lobule (LIP), precuneus (LPC), and inferior temporal (LIT) cortex responded more to nouns. MVPA revealed a double-dissociation in semantic sensitivity: classification was more accurate among verbs than nouns in the LMTG, and among nouns than verbs in the LIP, LPC, and LIT. However, classification was similar for verbs and nouns in the LIFG, and above chance for the non-preferred category in all regions. These results suggest that the meanings of verbs and nouns are represented in partially non-overlapping networks.

## Introduction

Words separate into classes based on the types of meanings they typically convey and the grammatical role they play in sentences. A key distinction that occurs across languages is the one between nouns and verbs (Sapir, 1921). Nouns tend to refer to entities (e.g. “the swan”, “the barn”), whereas verbs tend to describe events and relations among entities (e.g. “to lick”, “to sparkle”), thus reflecting the basic propositional acts of reference and predication (Langacker 1987, 2008). Within the structure of sentences, nouns serve as arguments and verbs as predicates (Bhat 2000; Croft 2005; Peterson 2007; Sasse 1993).

Because of its common occurrence across languages and its primacy within them, the noun verb distinction is a candidate organizing principle of the mental lexicon and its neural basis (Caramazza and Hillis 1991; Hillis and Caramazza 1995). Early neuropsychological studies showed that grammatical category affects the neural representations of words: focal brain damage leads to a disproportionate impairments with verbs in a subset of patients, while others are more impaired in processing nouns (Goodglass et al. 1966; Luria and Tsvetkova 1967; Miceli et al. 1984, 1988; McCarthy and Warrington 1985; Zingeser and Berndt 1990; Caramazza and Hillis 1991; Damasio and Tranel 1993; Daniele et al. 1994; Shapiro et al. 2000; Collina et al. 2001; Rapp and Caramazza 2002; Luzzatti et al. 2002; Shapiro and Caramazza 2003a, 2003b; Laiacona and Caramazza 2004; Aggujaro et al. 2006). For example, when asked to describe a scene, a subset of patients have difficulty naming the events depicted and fail to produce verbs for specific actions, either omitting the verbs altogether or using general verbs such as “be” or “do.” By contrast, patients with selective deficits for nouns have difficulties in naming the objects in the scene and tend to use generic terms such as “things”, “stuff” (Mätzig et al. 2009).

Consistent with the dissociations documented in the neuropsychological literature, neuroimaging studies with healthy participants have identified cortical regions that are preferentially recruited during either verb or noun processing (for reviews see Vigliocco et al. 2011; Crepaldi et al. 2013). Two regions that emerge across studies as specifically relevant to verb processing are the left middle temporal gyrus (LMTG) and the left inferior frontal gyrus (LIFG). Both regions have been found to respond more to verbs than nouns, adjectives and non-linguistic stimuli, across a variety of tasks, including semantic similarity judgments, lexical decision, and synonym judgments (Martin et al. 1995; Perani et al. 1999; Fujimaki et al. 1999; Kable et al. 2002, 2005; Li et al. 2004; Tyler et al. 2003, 2008; Davis et al. 2004; Tranel et al. 2005; Bedny and Thompson-Schill 2006; Thompson et al. 2007; Kemmerer et al. 2008; Liljeström et al. 2008; Bedny et al. 2008, 2011, 2014; Yu et al. 2011, 2012). Responses in these regions are observed not only to action verbs that involve motion (e.g. “to stroke” and “to give”), but also mental state (e.g. “to think” and “to want”) and other verb types (e.g. verbs of emission, such as “to glow”, change of state verbs, such as “to rust”) (Grossman et al. 2002; Davis et al. 2004; Bedny and Thompson-Schill 2006; Kemmerer et al. 2008; Bedny et al. 2008, 2011, 2014). A separate set of cortical areas has been identified as preferentially responsive to concrete nouns, including the left inferior parietal lobule (LIP) and angular gyrus, the left inferior temporal cortex (LIT), and the precuneus (PC) (Fujimaki et al. 1999; Li et al. 2004; Marangolo et al. 2006; Thompson et al. 2007; Liljeström et al. 2008; Berlingeri et al. 2008; Tyler et al. 2004; Shapiro et al. 2005, 2006; Bedny and Thompson-Schill 2006). Together these findings suggest that verbs and nouns are processed by partially non-overlapping cortical networks.

What role do these verb- and noun-responsive cortical areas play in lexical processing? One possibility is that brain regions responding preferentially to one grammatical class are selectively involved in representing lexical-semantic information that is associated with that particular grammatical category. For example, the LMTG might preferentially represent information about the meanings of verbs, whereas noun-responsive regions might be preferentially involved in representing the meanings of concrete nouns. Such a division of labor could occur in part because the dimensions that capture the meanings of verbs are partially distinct from those that are relevant to the meanings of nouns.

As noted above, verbs refer to events that are situated in time, whereas concrete nouns tend to refer to entities, which are fixed in time (Garbin et al. 2012; Bedny et al. 2014; Lapinskaya et al. 2016). Thus, the semantic dimensions distinguishing among verbs include whether they involve change over time (e.g. “to stroke”) or an ongoing state (e.g. “to want”) and the internal causal structure of the events to which they refer, such as its ultimate result (e.g. break, cover) or its manner (e.g. hit, shake) (Levin 1993; Levin and Hovav 1995). Since verbs relate different entities to each other, their meanings include a relational structure (Frawley 1992; Jackendoff 1983). For instance, part of the distinction between the verbs “to chase” and “to run” is that the former requires two arguments, namely a chaser and an agent/object being chased, while the latter only requires one argument. These aspects of verb semantics have implications for the verb’s grammatical behavior. Therefore, “the dog chased the cat” is more felicitous than “the dog chased”, but it is perfectly acceptable to say, “the dog ran”. One hypothesis is that verb-responsive areas, such as the LMTG, are involved in representing temporal and relational semantic information that is particularly relevant to verbs. By contrast, noun-responsive areas are involved in representing information relevant to the meaning of object-nouns, such as animacy, function or shape (Kemmerer 2017). In this regard, the dissociation between noun and verb areas may be akin to distinctions between higher-order visual regions in IT cortex that are preferentially involved in processing different types of objects (e.g. faces, scenes or body parts) and code visual dimensions that are most relevant to their preferred object category (e.g. face responsive areas coding nose shape, distance between the eyes) (Kanwisher et al. 1997; Epstein and Kanwisher 1998; Downing et al. 2001; Tsao and Livingston 2008).

However, multiple alternative possibilities regarding the cognitive role of verb- and noun-responsive regions remain open. One possibility is that all or some of the preferential responses to one grammatical class reflect different processing demands imposed by verbs and nouns on brain regions that are not preferentially involved in noun or verb representation *per se* (Siri et al. 2008). For example, regions that respond more to verbs than nouns could do so because they are involved in morphological processing, since verbs have a richer morphology in English (Tyler et al., 2001, 2003, 2004, 2008). Alternatively, brain regions generally involved in lexical retrieval might respond more to verbs than nouns because the former are more difficult to retrieve (Cordier et al. 2013; Bogka et al. 2003; Szekely et al. 2005; Engelkamp et al. 1990; Earles and Kersten 2000). Relative to nouns, verbs tend to be more context-dependent and have a wider range of meaning interpretations (Gentner 1981), which could make a single meaning more difficult to retrieve (Thompson-Schill et al. 1997, 1998). In particular, it has been suggested that inferior frontal responses to verbs are related to retrieval or morphosyntactic processing demands (Thompson-Schill et al. 2005; Tyler 2001, 2004).

The goal of the current study was to test the hypothesis that a subset of verb- and noun-responsive cortical regions are involved in representing the meanings of verbs and nouns. Based on prior work we hypothesized that the LMTG is preferentially involved in encoding verb meanings, whereas the IP, Precuneus (PC) and inferior temporal cortex (IT) are preferential involve in representing the meanings of object nouns. We predicted that these regions would be sensitive to semantic distinctions among verbs and among nouns. Furthermore, we predict a double-dissociation in multi-voxel sensitivity: verb-responsive (LMTG) would be more sensitive to distinctions among verbs whereas noun-responsive regions (LIP, LPC, LIT) would be more sensitive to distinctions among nouns. Since previous studies have suggested that the LIFG is active during verb processing because of morphosyntactic or retrieval demands, we did not predict preferential decoding among verbs (or nouns) in this region.

To test these predictions, we used multi-voxel pattern analysis (MVPA) to measure the spatial population code within noun and verb responsive cortical areas (Haxby et al. 2014). Several prior studies have shown that the spatial patterns of activation within temporal, parietal and prefrontal cortices, as measured by MVPA, are sensitive to semantic distinctions among words. Most studies thus far have focused on distinctions among concrete object nouns (Fairhall and Caramazza 2013; Wang et al. 2013; Simanova et al. 2014; Correia et al. 2014). For example, Fairhall and Caramazza (2013) showed that regions within the left temporal lobe are sensitive to distinctions among nouns referring to fruit, tools, clothes mammals and birds (see also Simanova et al. 2014, and Kumar et al. 2017 for similar design with place nouns). As for verbs, one study found that in the angular gyrus (AG), patterns of activation for two-word phrases were more similar when these phrases shared the same verb (Boylan et al. 2015). Another recent study found that fronto-temporal patterns of activation across action-verbs and object-nouns are correlated with their semantic similarity structure, as measured by latent semantic analysis (LSA) (Carota et al. 2017). Whether verb and noun responsive areas show sensitivity to semantics and whether they do so differentially across grammatical class has not yet been tested.

To test this prediction, we first localized brain regions responding more to verbs (LMTG and LIFG) and more to nouns (LIP, LPC, LIT) in each individual participant using a previously established paradigm (Bedny et al. 2008, 2014). Then, we tested whether the spatial patterns of activation within verb- and noun-responsive regions are more similar for verbs and nouns belonging to the same as opposed to different semantic classes – for instance, hand-action verbs (e.g. “to prod” and “to stroke”) should be more similar to each other than to light emission verbs (e.g. “to glow” and “to sparkle”).

Verb and noun stimuli were chosen from two broad semantic classes (verbs: actions vs. emission; nouns: animals vs. places), each including two narrower subclasses (action verbs: mouth actions, e.g. “to chew”, and hands actions, e.g. “to stroke”; emission verbs: sound, e.g. “to clang”, and light, e.g. “to sparkle”; animal nouns: birds, e.g. “the sparrow”, and mammals, e.g. “the fox”; place nouns: manmade, e.g. “the igloo”, and natural, e.g. “the meadow”). A secondary question of interest was whether verb/noun preferring regions would be sensitive to broad semantic distinctions only (e.g. action vs. emission) or also to the narrow subclasses (e.g. hand vs. mouth actions). We might expect that cortical areas that are sensitive to semantic information should show more pronounced distinctions for the broader classes but should also distinguish between the fine-grained semantic categories. On the other hand, some accounts posit that semantic information of different granularity is represented in different neural networks, particularly in the case of verbs (Kemmerer, two-part theory of verb meanings).

## Materials and Methods

### Participants

Thirteen individuals participated in the study (9 women, age range 19–56, mean age=34, SD=10). All participants were native English speakers and no history of neurological conditions (screened through self-report). Informed consent was obtained in accordance with the Johns Hopkins Medicine Institutional Review Boards.

### Stimuli and Procedure

Participants heard pairs of words and judged how related in meaning they were on a scale from 1 “*not at all similar*” to 4 “*very similar*”. Word stimuli consisted of 144 words, 18 words in each of the following 8 semantic categories: light emission verbs (e.g., “*to glow*” – henceforth, light verbs for brevity), sound emission verbs (e.g. “*to boom*” – henceforth, sound verbs), hand-related action verbs (e.g. “*to stroke*” – henceforth, hand verbs), mouth-related action verbs (e.g. “*to bite*” – henceforth, mouth verbs), bird nouns (e.g. “*the sparrow*”), mammal nouns (e.g. “*the deer*”), manmade place nouns (e.g. “*the dungeon*”), and natural place nouns (e.g. “*the creek*”); All verb stimuli occurred in the infinitive and all noun stimuli were preceded by the article “the”, to mark grammatical category (See Supplementary Material, Appendix 1).

Words were matched across semantic categories in syllable length based on CMU Pronouncing Dictionary (Weide 1998) (Table 1; One-way ANOVA F_(7, 136)_=0.38, p=0.91). Words were also matched in familiarity, based on ratings collected using Amazon Mechanical Turk (AMT)(One-way ANOVA F_(7, 136)_=1.07, p=0.38). Participants (N=15, 5 women; mean age=27, SD 7, all English native speaker) rated how often they had encountered each word on a scale from 1 “*never*” to 7 “*very often*”. Since words were highly familiar, but infrequent (e.g. “*to beep*” familiarity=4.67, log10(freq.)=0.34), we chose to match the words on familiarity because this variable has previously been shown to be a better predictor of performance (e.g. Connine et al. 1990).

**Table 1.** Word stimuli

|  | Familiarity | Syllable length | Log10 Frequency |
| --- | --- | --- | --- |
| **Verbs** | **3.66 (0.69)** | **1.31 (0.44)** | **0.56 (0.46)** |
| Light | 3.72 (0.68) | 1.39 (0.5) | 0.67 (0.41) |
| Sound | 3.63 (0.76) | 1.25 (0.39) | 0.54 (0.47) |
| Hand | 3.4 (0.6) | 1.25 (0.43) | 0.44 (0.47) |
| Mouth | 3.89 (0.69) | 1.33 (0.45) | 0.59 (0.5) |
| **Nouns** | **3.6 (0.53)** | **1.33 (0.46)** | **0.85 (0.49)** |
| Birds | 3.52 (0.47) | 1.39 (0.5) | 0.68 (0.39) |
| Mammals | 3.64 (0.46) | 1.25 (0.43) | 0.82 (0.53) |
| Manmade places | 3.53 (0.75) | 1.28 (0.43) | 0.75 (0.57) |
| Natural places | 3.7 (0.37) | 1.39 (0.5) | 1.16 (0.34) |

Means and standard deviations (presented in parentheses next to the mean) of familiarity, syllable length and frequency for all word categories.
**Log10(freq.) statistics:** One-way ANOVA, $F_{(7, 136)}=3.92$ , $p<0.001$ . Word frequency information gathered from the Corpus of Contemporary American English (COCA – Davies, 2008).

Words from each category were divided into two non-overlapping sets (9 words per category per set) to be used separately in even/odd runs. Within each set, we generated all possible within category pairs (36 pairs per category per set). Word pairs were presented in blocks of four. All blocks contained only words from a single semantic category and each word occurred only once per block (e.g. light verbs block: “*to shine – to glisten*”, “*to flash – to gleam*”, “*to twinkle – to glare*”, “*to glimmer – to blaze*”). Blocks were 16s long, separated by 10s of rest. Each trial within a block was 4s long, including two words (0.9s each), an inter-word interval 0.25s and the response period (1.95s).

There was a total of 144 blocks (18 per word category) divided evenly into 8 runs. Blocks in even/odd runs contained stimuli exclusively from one of the two sets, ensuring that words were not repeated across even/odd runs (e.g. “*to flash*” occurred only in even runs). Word pairs, blocks and runs were presented in one of two pseudo-random orders, counterbalanced across participants. For every run, each word category was distributed evenly throughout the run to minimize position effects.

A female native English speaker recorded the word stimuli. The audio files were normalized to each other in volume with respect to root-mean square (RMS) amplitude and adjusted to have equal durations (0.9s). The stimuli were presented over MRI-compatible earphones at the maximum comfortable volume for each participant. Participants indicated their responses by using an MRI-compatible button pad. Because data from the current participants were also used as a control group for a separate study with blind individuals, participants wore a light exclusion blindfold throughout the experiment.

### fMRI Data Acquisition

MRI structural and functional data of the whole brain were collected on a 3 Tesla Phillips scanner using a 32 channel head coil. T1-weighted structural images were collected using an MPRAGE pulse sequence in 150 axial slices with 1mm isotropic voxels. Functional, BOLD images were collected in 36 axial slices with 2.4 × 2.4 × 3 mm voxels and TR=2s.

### fMRI Data Analysis

#### Preprocessing

Data analyses were performed using FSL, Freesurfer, the Human Connectome Project workbench, and custom software (Dale, Fischl, and Sereno 1999; Glasser et al. 2013; Smith et al. 2004). Functional data were corrected for subject motion using FSL’s MCFLIRT algorithm (Jenkinson et al. 2002), high pass filtered to remove signal fluctuations at frequencies longer than 128 seconds/cycle, and then resampled to a cortical surface model and smoothed with a 2mm FWHM Gaussian kernel to regularize the data on the cortical surface. All subsequent analyses were surface-based. Subject-specific cortical surface models were generated using the automated Freesurfer pipeline and visually inspected to assure accuracy.

#### Whole-brain univariate analysis

The functional data were spatially smoothed with a 6mm FWHM Gaussian kernel and prewhitened to remove temporal autocorrelation. Each of the verb and noun categories were entered as a separate predictor in a general linear model (GLM) after convolving with a canonical hemodynamic response function. We also included the first derivative as a covariate of no interest. Each run was modeled separately, and runs were combined within subject using a fixed-effects model. Group-level random-effects analyses were corrected for multiple comparisons at vertex level with p < 0.05 threshold false discovery rate (FDR) across the whole cortex (Genovese et al. 2002). Additionally, a nonparametric permutation test was used to cluster-correct at p < 0.01 family-wise error rate (FWER).

#### ROIs definition

We defined individual subject functional ROIs to be used in the MVPA analysis. For each subject, we defined two verb-responsive ROIs and four noun-responsive ROIs: verb ROIs left middle temporal gyrus (LMTG), the left inferior frontal gyrus (LIFG); noun-responsive ROIs left inferior parietal cortex (LIP), left inferior temporal cortex, both laterally (latLIT) and medially (medLIT), and left precuneus (PC). These anatomical regions were chosen because they have been observed to respond preferentially to verbs or nouns in previous studies, and showed verb/noun preferences in the univariate analysis in the current study (Crepaldi et al. 2013 for a review).

We employed a two-step procedure to identify subject-specific functional ROIs. First, whole-brain results for the Verbs>Nouns contrast were used to identify group search spaces in the anatomical location of LMTG, LIFG, LIT, left PC, left latIT and medIT. Next individual functional ROIs were defined for each subject by taking the top 300 active vertices for the Verbs>Nouns and Nouns>Verbs contrasts, for verb-responsive and noun-responsive ROIs respectively. Note that although the focus of the current paper was on MVPA patterns within these ROIs, we also report the univariate signal for each verb/noun category in the Supplementary Materials.

#### MVPA ROI analysis

MVPA was used to test whether verb- and noun-responsive regions are more sensitive to lexico-semantic differences among verbs and among nouns, respectively, using PyMVPA toolbox (Hanke et al. 2009). For each ROI in each participant, a linear support vector machine (SVM) classifier was used to separately decode among the four verb categories and among the four noun categories. In each vertex in each participant’s ROIs, we obtained one sample labeled by semantic category per block by averaging BOLD signal across time points over a block duration (16s). Time points for each block were defined as block onset plus 4s delay to account for the hemodynamic lag. Each vertex’s block sample was then normalized (z-scored) with respect to the mean signal for the vertex across the task blocks during a given run, such that the mean of each vertex across the run was set to 0 and standard deviation to 1. Normalization was applied to data from verbs and nouns separately, removing differences in mean signal across grammatical category. The classifier was then trained on half of the data (e.g. even runs) and tested on the other half (e.g. odd runs). Classification accuracy was then averaged across the two training/test splits. Importantly, individual verbs and nouns did not repeat across even and odd runs. Thus, the classifier was trained on one subset of words and tested on a different subset.

We compared classifier performance within each ROI to chance (25%) and across verbs and nouns. We also tested for an interaction between grammatical class (verbs vs. nouns) and ROI. Significance was evaluated against an empirically generated null distribution using a combined permutation and bootstrap approach (Stelzer, Chen and Turner 2013; Schreiber and Krekelberg 2013). In this approach, t- and F-statistics obtained for the observed data are compared against an empirically generated null distribution. We report the t- and F-values obtained for the observed data and the non-parametric p-values, where p corresponds to the proportion of the shuffled analyses that generated a comparable or higher t/F value.

The null distribution was generated using a balanced block permutation test by shuffling the block labels within run 1000 times for each subject (Schreiber and Krekelberg 2013). Then, a bootstrapping procedure was used to generate an empirical null distribution for each statistical test across participants by sampling one permuted accuracy value from each participant’s null distribution 15,000 times (with replacement) and running each statistical test on these permuted samples, thus generating a null distribution of 15,000 statistical values for each test (Stelzer, Chen and Turner 2013).

Confusion matrices were generated to describe how well the classifier performed on each pairwise distinction among verbs and nouns (e.g. manmade places vs. natural places). For any pair of categories or subcategories, the confusion matrix yields two measures of classification performance: the percentage of correctly classified trials, or hit rate (H), and the percentage of misclassified trials, or false alarm rate (F) (Haxby et al. 2014). Classification and misclassification frequencies can be compared using a signal detection theory framework (Swets et al. 1961; Green and Swets 1966). Within each ROI, we computed A’, a non-parametric estimate of discriminability 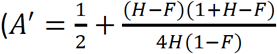; Pollack and Norman 1964; Grier 1971; Stanislaw and Todorov 1999), to assess the classifier’s ability to distinguish (1) between “major” semantic categories (i.e. animal vs. place nouns, action vs. emission verbs), and (2) between the semantic subcategories (i.e. birds vs. mammals, manmade vs. natural places; mouth vs. hand actions, sound vs. light emission verbs). An A’ of 0.5 corresponds to chance performance, while 1.0 indicates perfect discriminability. Paired student t-tests were used to compare A’ values.

#### Whole-brain searchlight MVPA analysis

A whole-brain SVM classifier was used to decode independently among verbs and among nouns over the whole cortex using a 6mm radius circular searchlight (according to geodesic distance, to better respect cortical anatomy over Euclidean distance) (Glasser et al. 2013). This yielded for each participant two classification maps (one for verbs and one for nouns) indicating the classifier’s accuracy in a neighborhood surrounding every vertex. Individual subject searchlight accuracy maps were averaged, and this group-wise map was thresholded using PyMVPA implementation of the two-step cluster-thresholding procedure described in Seltzer et al. 2013 (Hanke et al. 2009). This procedure permutes block labels within participant and run to generate a null distribution within subject (100 times) and then samples from these (10,000) to generate a group-wise null distribution (as in the ROI analysis.) The whole-brain searchlight maps are then thresholded using a combination of vertex-wise threshold (p<0.001 uncorrected) and cluster size threshold (FWER p<0.05, corrected for multiple comparisons across the entire cortical surface).

## Results

### Behavioral results

Participants judged verbs and nouns to be equally similar in their meaning (verbs: mean=2.08, SD=0.33; nouns: mean=2.03, SD=0.33; paired t-test t_(12)_=0.76, p=0.46). Among verbs, light verbs were judged to be more similar than any other verb category (light mean=2.74, SD=0.6, sound mean=1.8, SD=0.29, hand mean=1.86; SD=0.31; mouth: mean=1.9, SD=0.26, Repeated measures ANOVA F_(3,36)_=43.15, p<0.001). Among nouns, manmade places were judged to be less similar than the other categories (birds mean= 2.28, SD=0.52, mammals mean=2.19, SD=0.44, manmade places mean=1.6, SD=0.13, natural places mean=2.05, SD=0.39, Repeated measures ANOVA F_(3,36)_=17;68, p<0.001); Participants’ reaction times were not different for verbs versus nouns (verbs mean=1.57s, SD=0.16; nouns mean=1.59s, SD=0.17; paired t-test t_(12)_=1.45, p=0.17). Among verbs, responses were slower for hand verbs than the other verbs (light mean=1.57, SD=0.21, sound mean=1.58, SD=0.16, hand mean=1.61; SD=0.17; mouth: mean=1.52, SD=0.16, Repeated measures ANOVA F_(3,36)_=3.69, p=0.02). Among nouns, responses were slower for birds and natural places than for mammals and manmade places (birds mean=1.60, SD=0.22, mammals mean=1.56, SD=0.17, manmade places mean=1.57, SD=0.16, natural places mean=1.64, SD=0.19, Repeated measures ANOVA F_(3,36)_=3.61, p=0.02).

### fMRI Results

#### Verb and noun preferring cortical networks identified in mean signal

Greater activation for verbs than for nouns was found bilaterally in the MTG, a portion of the LIFG, as well as the lateral occipital cortex (LOC) (p<0.01 FWER, Figure 1 A). In contrast, brain activity was greater for nouns than verbs bilaterally in an IP region, in the superior and middle frontal gyri, in the left IT cortex both laterally (LlatLIT) and medially (LmedLIT), and medially in PC. The extent of activation for verbs and nouns was larger in the left hemisphere (Table 2 and 3).

**Figure 1:**
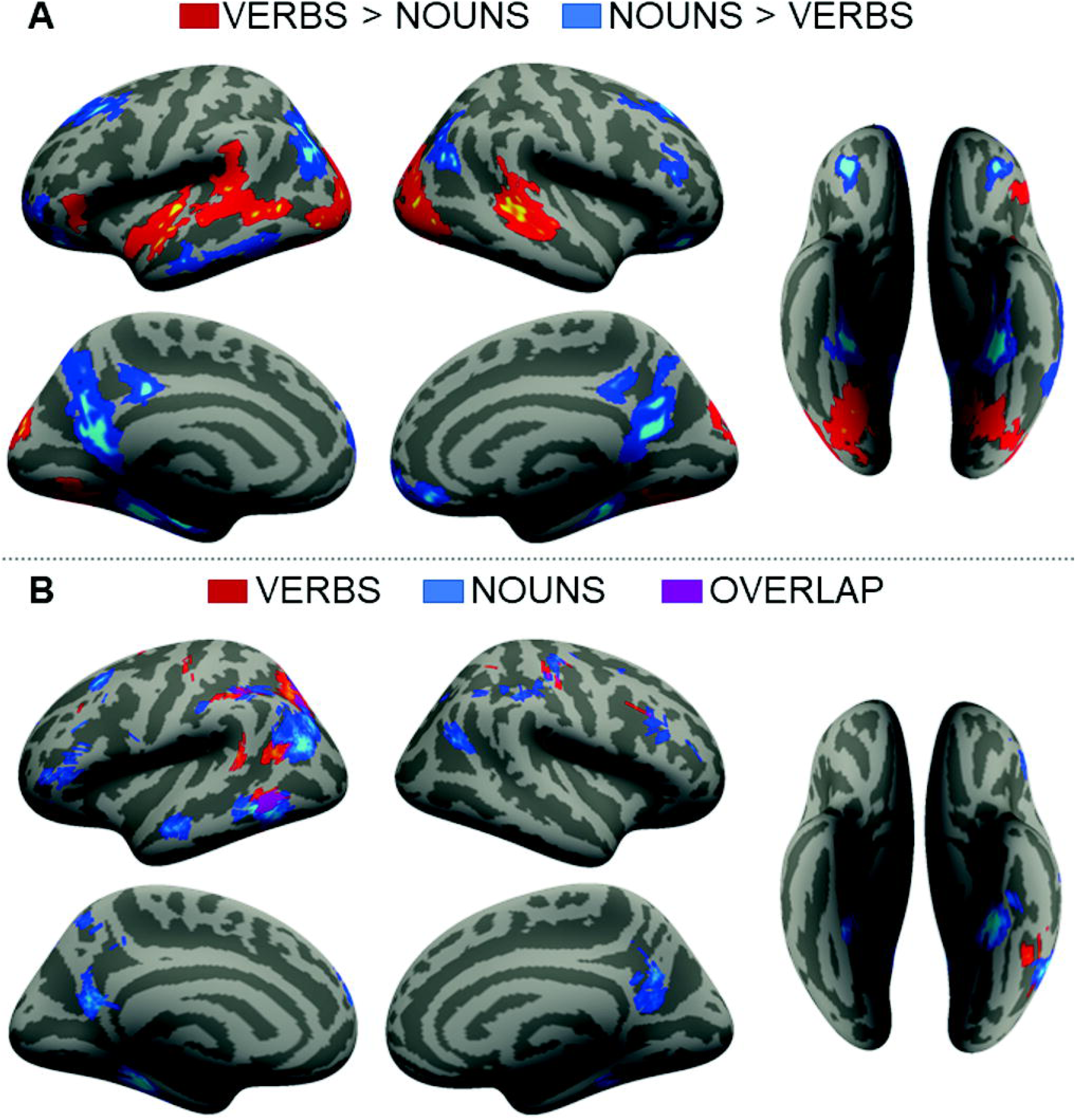
**A)** whole-brain group maps, p<0.01 FWER. Red: verbs, blue: nouns. **B)** MVPA searchlight group maps, vertex-wise accuracy significance p<0.001, cluster-thresholded FWER p<0.05. Red: verbs, blue: nouns, purple: verbs/nouns overlap.

**Table 2.** Coordinates (MNI) and statistics for activation peaks produced by verbs>nouns (FWER 0.01)

| Brain Region | x | y | z | Peak t | mm<sup>2</sup> | P<sub>cluster</sub> |
| --- | --- | --- | --- | --- | --- | --- |
| Left hemisphere |  |  |  |  |  |  |
| Anterior Superior Temporal Gyrus | -43 | -23 | 0 | 11.23 | 1222.23 | 0.0028 |
|  | -59 | -5 | 0 | 7.78 |  |  |
| Posterior Middle Temporal Gyrus | -57 | -62 | 5 | 8.67 | 1584.51 | 0.0016 |
| Posterior Superior Temporal Gyrus | -63 | -35 | 12 | 6.87 |  |  |
| Lateral Occipital Cortex/Cuneus | -43 | -74 | -4 | 8.04 | 1666.5 | 0.0008 |
|  | -5 | -87 | 26 | 7.39 |  |  |
|  | -29 | -86 | 5 | 7.36 |  |  |
| Lingual/Fusiform Gyri | -14 | -70 | -5 | 6.43 | 813.66 | 0.0066 |
|  | -29 | -66 | -13 | 5.6 |  |  |
| Inferior Frontal Gyrus (pars triangularis) | -50 | 28 | 2 | 6.04 | 315.18 | 0.028 |
| Right hemisphere |  |  |  |  |  |  |
| Posterior Superior Temporal Sulcus and Gyrus | 52 | -33 | 1 | 9.19 | 1236.6 | 0.0016 |
| Lateral Occipital Cortex | 41 | -83 | 4 | 7.95 | 1817.45 | 0.0006 |
|  | 34 | -85 | 15 | 6.7 |  |  |
|  | 4 | -81 | 31 | 5.43 |  |  |
| Fusiform gyrus | 28 | -68 | -10 | 6.61 | 769.34 | 0.0038 |

**Table 3.**
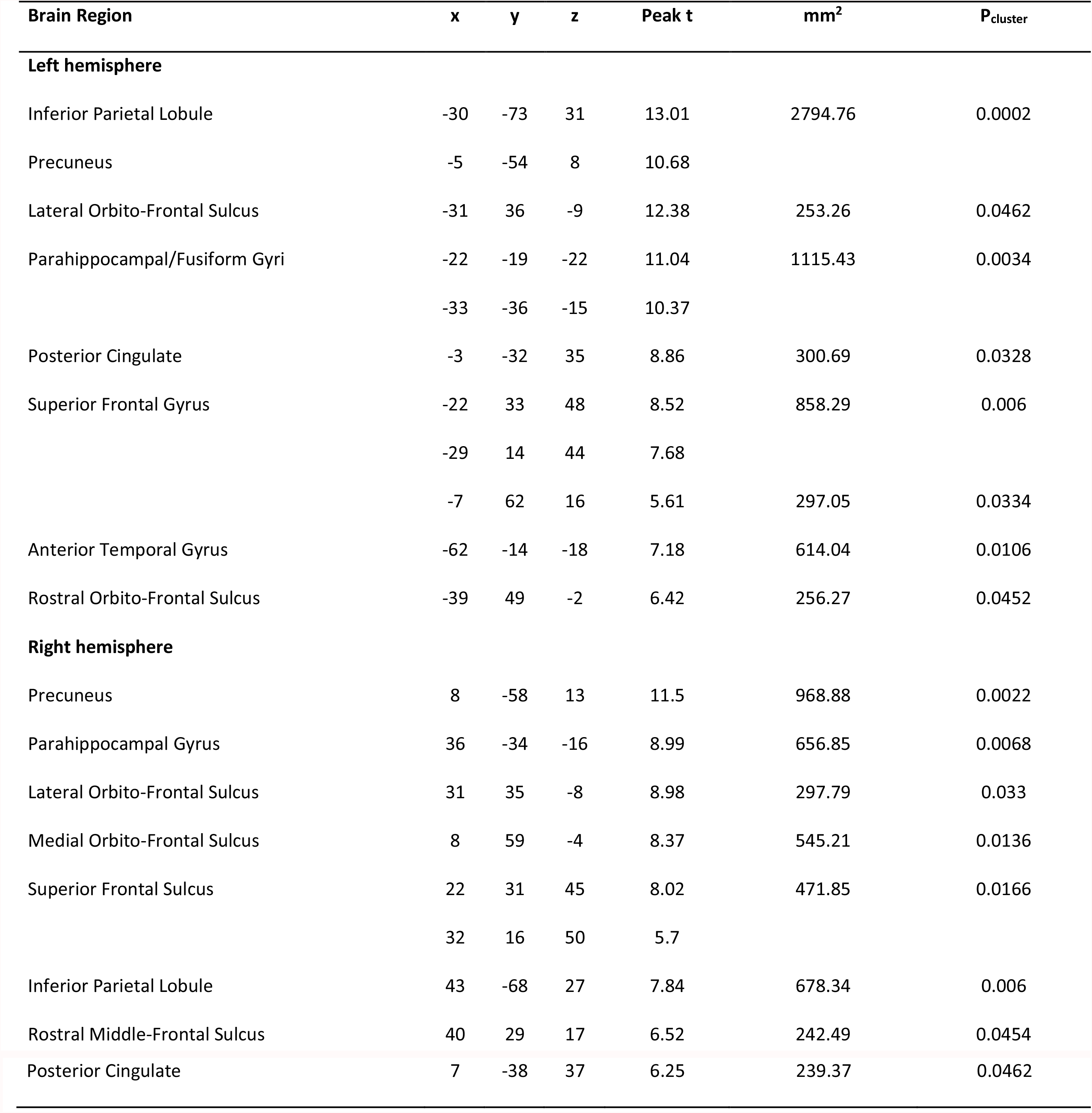
Coordinates (MNI) and statistics for activation peaks produced by nouns>verbs (FWER 0.01)

#### MVPA distinctions among verb and among noun types within verb- and noun-responsive ROIs

Classification was tested within three verb-responsive (LMTG and LIFG, LLOC) and four noun-responsive (LIP, LPC, LmedIT and LlatIT) ROIs within the left hemisphere. Classifier performance for both verbs and nouns was significantly above chance (25%) in all ROIs, except the occipital cortex (LLOC, verbs: 26%, t_(12)_=0.62, p=0.28; nouns: 29%, t_(12)_=1.29, p=0.11): LMTG (verbs: 36%, t_(12)_=3.51, p<0.001; nouns: 30%, t_(12)_=2.73, p<0.005), LIFG (verbs: 30%, t_(12)_=2.82, p<0.01; nouns: 34%, t_(12)_=2.78, p<0.01), LIP (verbs: 34%, t_(12)_=4.3, p<0.000; nouns: 43%, t_(12)_=7.08, p<0.000), LPC (verbs: 29%, t_(12)_=2.02, p<0.05; nouns: 38%, t_(12)_=4.88, p<0.000), and LlatIT (verbs: 31%, t_(12)_=4.45, p<0.000; nouns: 41%, t_(12)_=5.77, p<0.000). In the LmedIT, classification was significantly above chance for nouns only (verbs: 28%, t_(12)_=1.3, p=0.09; nouns: 40%, t_(12)_=5.11, p<0.000)(Figure 2).

**Figure 2:**
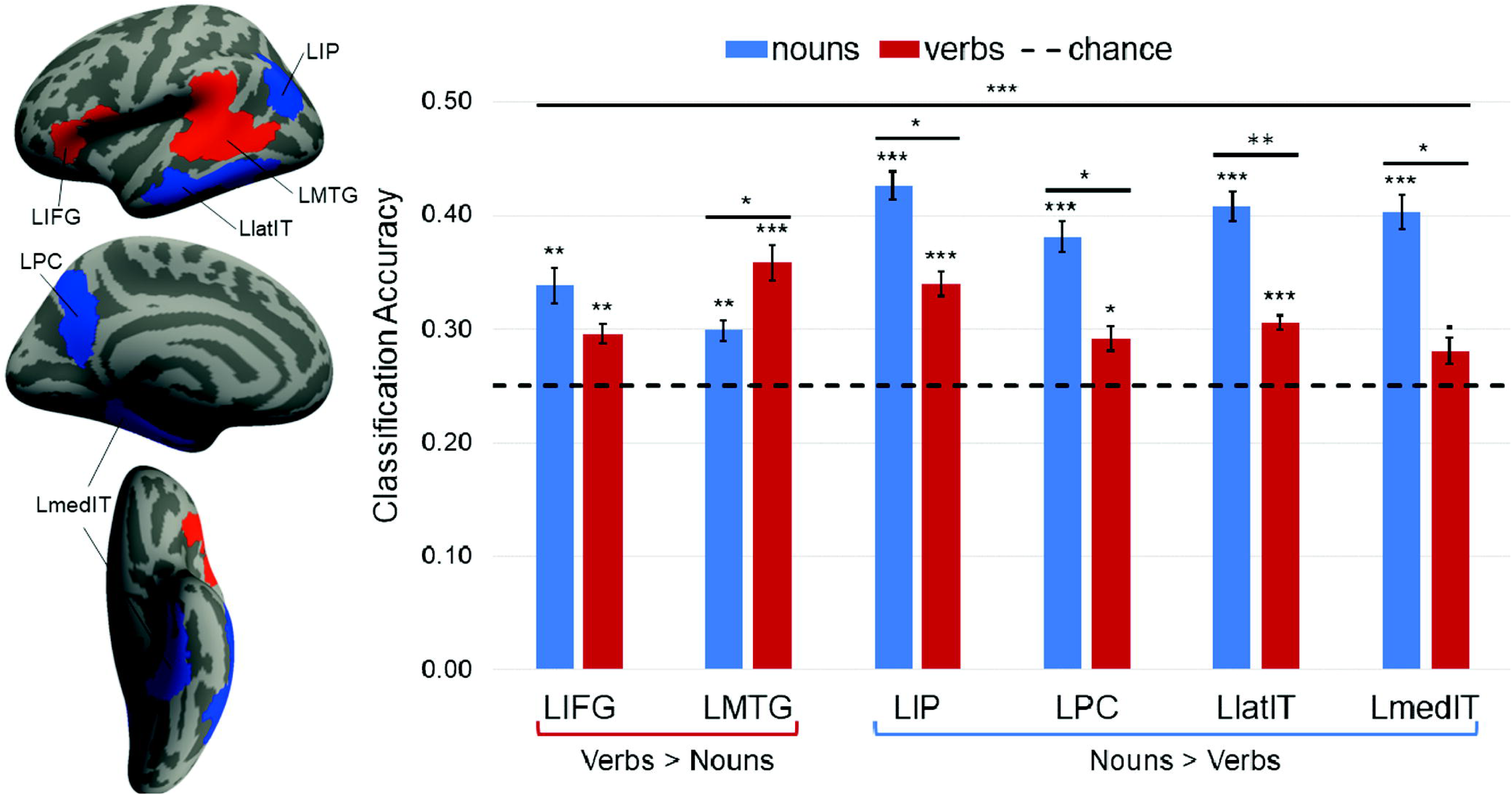
Group search spaces used to define individual subject functional ROIs in the left hemisphere. Classifier accuracy in verb (LIFG and LMTG) and noun (LIP, LPC, LlatIT and LmedIT) selective regions. Chance: 25%. Signif. codes: 0 ‘***’ 0.001 ‘**’ 0.01 ‘*’ 0.05 ‘.’ 0.1 ‘’ 1

We observed a grammatical class (verbs, nouns) by ROI (LMTG, LIFG, LIP, LPC, LlatIT, LmedIT) interaction (Repeated measures ANOVA F_(5,60)_=7.06, p<0.000), a main effect of ROI (F_(5,60)_=3.29, p<0.05), and a main effect of grammatical class (F_(1,12)_=6.03, p<0.05)(Figure 2). The LMTG was more sensitive to distinctions among verbs than nouns (t_(12)_=2.11, p=0.05), whereas all noun-responsive regions (LIP, LPC, LlatIT and LmedIT) were more sensitive to the distinctions among nouns than verbs (LIP t_(12)_=2,67, p<0.05; LPC: t_(12)_=2.65, p<0.05; LlatIT t_(12)_=3.51, p<0.005; LmedIT: t_(12)_=2.85, p<0.05). By contrast, verbs and nouns were equally decodable in the LIFG (t_(12)_=1.12, p=0.29).

Inspection of the confusion matrices revealed that in verb-responsive LMTG, the classifier successfully discriminated between the “major” verb semantic categories of action and emission verbs (A’=0.67, t_(12)_=4.46, p<0.000), as well as between the smaller subclasses, albeit to a lesser degree (mouth vs. hand actions A’=0.63, t_(12)_=2.43, p<0.05; sound vs. light emission A’=0.59, t_(12)_=1.62, p=0.06) (Figure 3). In the verb-responsive LIFG, the only significant discrimination among verbs was between sound and light emission verbs (A’=0.66, t_(12)_=3.5, p<0.01). The classifier also successfully discriminated between the “major”, but not the “minor”, noun semantic categories in both verbs selective regions (LMTG: A’=0.61, t_(12)_=2.45, p<0.05; LIFG: A’=0.64, t_(12)_=2.79, p<0.01).

**Figure 3:**
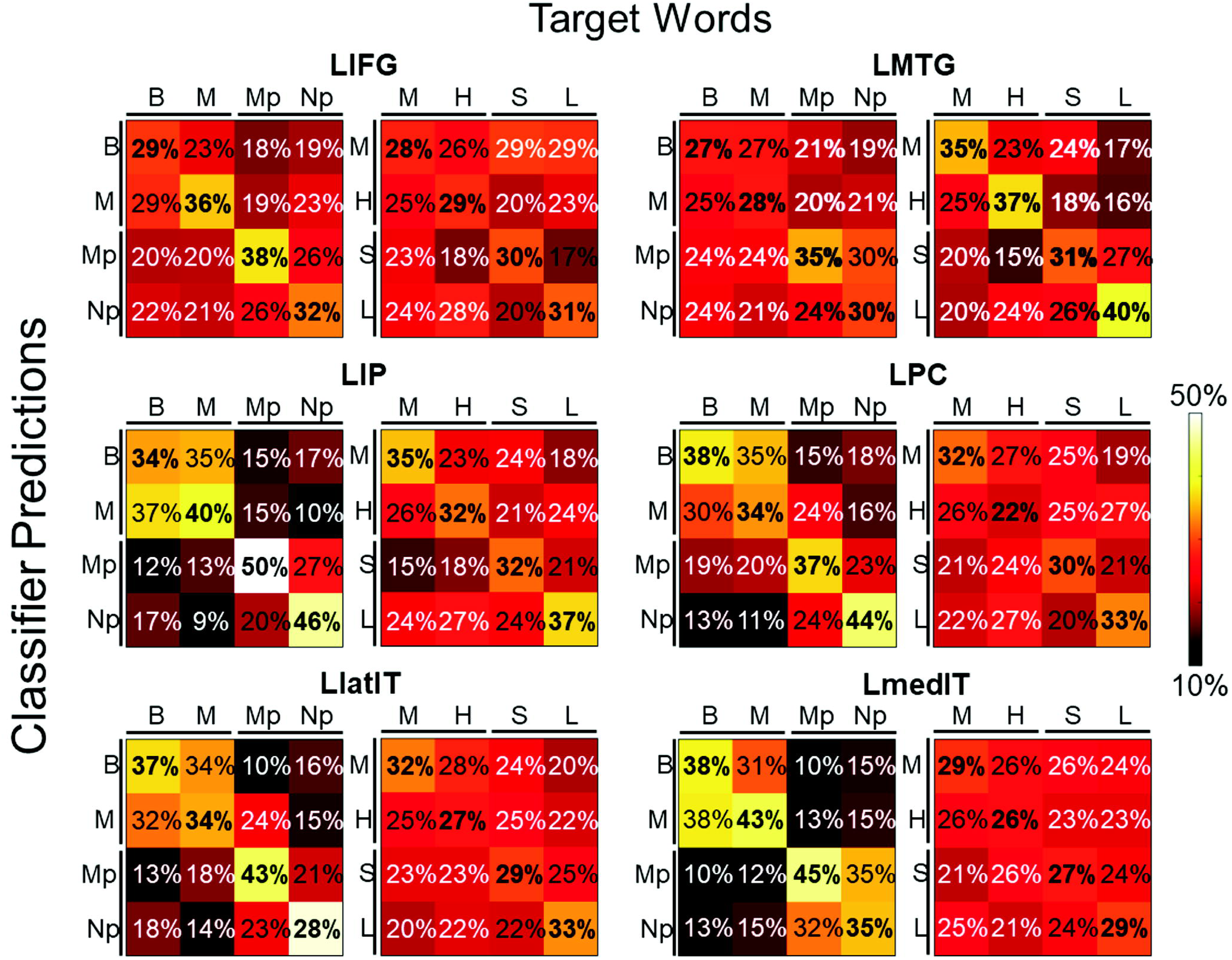
Confusion matrices for verbs- and nouns-selective ROIs providing the percentage of predicted correct classifications (diagonals) and misclassifications (off diagonals). Labels, nouns: B: birds, M: mammals, Mp: manmade places, Np: natural places; verbs: M: mouth, H: hand, S: sound, L: light.

In all the noun-responsive regions (LIP, LPC, LlatIT and LmedIT), the classifier successfully discriminated between the “major” noun semantic categories of animals and places (LIP: A’=0.80, t_(12)_=11.74, p<0.000; LPC: A’=0.72, t_(12)_=4.19, p<0.001; LlatIT: A’=0.75, t_(12)_=7.76, p<0.000; LmedIT: A’=0.82, t_(12)_=15.06, p<0.000). Classification was also above chance for manmade versus natural places in all but one noun-responsive ROI (LIP: A’=0.72, t_(12)_=4.97, p<0.001; LPC: A’=0.68, t_(12)_=3.5, p<0.005; LlatIT: A’=0.71, t_(12)_=4.8, p<0.001, LmedIT: A’=0.54; t_(12)_=0.99, p=0.17). By contrast, birds and mammals were not distinguishable in any ROI (LIP: A’=0.49, t_(12)_=-0.22, p=0.59; LPC: A’=0.51, t_(12)_=0.08, p=0.47; LlatIT: A’=0.52, t_(12)_=0.38, p=0.36, LmedIT: A’=0.54; t_(12)_=0.75, p=0.23).

In three of the four noun-responsive regions, the classifier also discriminated between the “major” verb semantic categories (LIP: A’=0.62, t_(12)_=4.07, p<0.001; trending in LPC: A’=0.55, t_(12)_=1.52, p=0.07; LlatIT: A’=0.59; t_(12)_=3.1, p<0.005, LmedIT: A’=0.54, t_(12)_=1.25, p=0.12). In the LIP, the classifier also discriminated successfully between mouth and hand actions (A’=0.61; t_(12)_=2.2, p<0.05), and sound and light emission verbs (A’=0.61; t_(12)_=1.81, p<0.05). In the LPC, the classifier successfully distinguished between light and sound emission verbs (A’=0.63; t_(12)_=1.94, p<0.05). No other classifications were above chance.

#### MVPA searchlight results

The searchlight analyses were broadly consistent with the ROI results (Figure 1 B). Classification among verbs was significantly better than chance in the posterior LMTG, although the searchlight classification peak was somewhat posterior and superior to the verbs>nouns peak observed in the univariate analysis. Verb classification was also above chance in LlatIT, as well as the posterior LIP. Classification among nouns was significantly better than chance in LIP, LmedIT and LlatIT (anterior and posterior), bilaterally in the PC and in the L IFG. Consistent with the univariate analysis and the ROI results, the verb and noun sensitive networks showed some overlap, but each had a distinctive neuroanatomical distribution.

## Discussion

Consistent with the idea that verb- and noun-responsive regions are involved in representing lexical semantic information, we find that the spatial patterns of activity within these areas are sensitive to distinctions among nouns and verbs. Furthermore, verb-responsive LMTG (but not LIFG) and all the regions showing elevated responses to nouns (LIP, LPC, LlatIT and LmedIT) are more sensitive to distinctions among their preferred grammatical class. In other words, we observed a double dissociation in the sensitivity of spatial patterns of activation to different lexical types. While the LMTG was more sensitive to distinctions among verbs, noun-responsive regions were more sensitive to distinctions among nouns. This suggests that the meanings of verbs and nouns are represented in partially non-overlapping neural systems.

### The left middle temporal gyrus (LMTG) is preferentially sensitive to the meanings of verbs

Verb-responsive LMTG was the only cortical region that was more sensitive to distinctions among verbs than among nouns. This finding is consistent with the large body of work showing preferential responses to verbs as compared to nouns in this area (Martin et al. 1995; Perani et al. 1999; Fujimaki et al. 1999; Kable et al. 2002, 2005; Li et al. 2004; Tyler et al. 2003, 2008; Davis et al. 2004; Tranel et al. 2005; Bedny and Thompson-Schill 2006; Thompson et al. 2007; Liljeström et al. 2008; Bedny et al. 2008, 2011, 2014; Yu et al. 2011, 2012). However, we also found that presence of a mean difference between verbs and nouns does not guarantee decoding among semantic types or a differential sensitivity for verbs as opposed to nouns. While the lateral occipital cortices were more responsive to verbs than nouns, neither verb nor noun distinctions could be decoded in occipital cortex. Conversely, a region within the LIFG that was more active during verb than noun processing was equally sensitive to distinctions among verbs and nouns showing, if anything, slightly more sensitivity to distinctions among nouns.

The present results suggest that the verb-responsive LMTG and LIFG play distinct roles in word processing. Only the LMTG appears to be specifically involved in verb-meaning representation. One possibility is that this LIFG supports general cognitive control functions involved in verb retrieval (Thompson-Schill et al. 2005). In English, verbs tend to be more semantically malleable and influenced by the sentence context than object nouns, potentially increasing the demand for selection among possible alternative meanings (Gentner 1981), and the LIFG is sensitive to this property of words, i.e. their semantic ambiguity (Grindrod et al. 2008; Rodd et al. 2005; Thompson-Schill et al. 1997, 1999; Bedny et al. 2007, 2008). Alternatively, the LIFG might contribute to morphosyntactic processing (Tyler 2001, 2004). Consistent with some previous proposals, the current results suggest that, unlike the LMTG, the LIFG is not specifically involved in representing verb as opposed to noun meanings.

What kind of verb-relevant information does the LMTG represent? One possibility is that the LMTG encodes the number and type of arguments that a verb takes, i.e. its argument-structure (Thompson et al. 2007; den Ouden et al. 2009; Hernandez et al. 2014). Consistent with this idea, we find that spatial patterns of activation within the LMTG are distinguishable when verbs have different numbers and types of arguments (e.g. divalent action verbs as opposed to monovalent emission verbs). For instance, the verb “to lick” requires two arguments, the agent doing the licking and the object being licked (e.g. “the dog licked the bone”), while emission verbs such as “to sparkle” are monovalent (e.g. “the diamond sparkled”) (Levin 1993; Levin and Hovav 1995). Action and emission verbs also differ in the type of arguments that can serve as their subjects. While the subjects of action verbs are intentional agents, those of emission verbs are most often forces/natural causes. The argument structure of verbs is intimately related not only to their meaning, but also to their grammatical behavioral in sentences. The present results are therefore consistent with the possibility that the LMTG and surrounding cortices play a role not only in semantic, but also in the grammatical processing of verbs.

However, a purely argument-structure or grammatical account of LMTG function cannot fully account for the available evidence. Previous studies suggest that LMTG responds not only to verbs, but also to event nouns (e.g. hurricane, rodeo). The LMTG response to event nouns is intermediate between its response to object nouns and verbs (Collina et al. 2001; Tabossi et al. 2010; Garbin et al. 2012; Bedny et al. 2014; Lapinskaya et al. 2016). The grammatical behavior of event nouns within sentences and their morpho-syntactic properties are akin to those of object nouns (“the concert” but not “concerting”). Semantically, however, event nouns resemble verbs in that their referents are situated in time, in addition to space (e.g. “during the concert”) (Langacker 1987, 2008). Its involvement in the processing of event nouns suggests that the LMTG plays a role in representing the meanings of words, above and beyond any role that it plays in grammatical processing.

Further support for the idea that the LMTG plays a role in semantic processing above and beyond any role in grammatical functions comes from the current study. We found that patterns of activity in the LMTG distinguish not only between verb classes that are grammatically different (to clap and to glow) but also between verbs that differ in subtler semantic dimensions (i.e. the effector of action hand e.g. to clap vs. mouth e.g. to bite and “manner” of emission light e.g. to glow vs. sound e.g. to boom); This finding suggests that the LMTG stores fine-grained representations of lexicalized events, including verbs and event nouns.

What is the format and content of LMTG semantic representations? Although this question remains largely to be address, several hints are available from prior studies. First, previous evidence suggests that the LMTG represents meanings in a modality-independent format. Although early hypotheses regarding LMTG function suggested imagistic visual motion representations (Martin et al. 1995), more recent evidence suggests that LMTG representations are neither visual nor specific to motion. Basic and biological motion perception tasks engage a network of areas that is neuroanatomically separate from the LMTG (Bedny et al. 2008; Bedny and Caramazza 2011). The LMTG responds just as much to abstract mental state verbs as it does not concrete action verbs (Grossman et al. 2002; Davis et al. 2004; Bedny and Thompson-Schill 2006; Kemmerer et al. 2008; Bedny et al. 2008, 2011, 2014). Finally, the LMTG is involved in verb comprehension even in individuals who have never experienced visual motion, i.e. those blind from birth (Bedny et al. 2011; Bedny and Caramazza 2011; Bedny and Saxe 2012). These lines of evidence demonstrate that verb meaning representations in the LMTG are modality-independent rather than “visual”, in that they do not depend on visual experience for acquisition, do not specifically represent information experienced through a particular sensory modality, and do not specifically represent perceptible information (e.g. motion) but rather any semantic information relevant to verbs. Together with prior evidence, the present results therefore suggest that the LMTG stores fine-grained modality-independent semantic representations of verbs and event nouns.

Many further questions remain open regarding LMTG’s contribution to semantic representations and processing. In future work, it will be important to uncover the specific dimensions of verb meaning and grammar that are represented in the LMTG and surrounding verb-responsive temporal cortex. Many candidate dimensions have been identified in linguistics, including whether the verb refers to a state (e.g. “to contain”) or an event (e.g. “to boil”), whether it describes an ongoing or a finished event (i.e. telicity), the agentivity of the verb’s subject (animate/inanimate) and whether the verb encodes the ultimate result of an event (e.g. break, cover) or its manner (e.g. hit, shake) (Levin and Hovav 1995). Future studies could use MVPA to uncover which of these dimensions are explicitly coded by neural populations. A further open question concerns the degree to which verb-responsive LMTG region is language specific. Previous studies suggest that recognition and categorization of action images and videos depends on distinct right-lateralized superior temporal sulcus (STS) and MTG regions (Vander Wyk et al. 2009; Pelphrey et al. 2004, 2005; Bedny et al. 2010). However, studies with matched verbal and non-verbal stimuli are needed to resolve this question. Finally, although the LMTG appears to contribute disproportionately to representations of verbs and event nouns as opposed to object nouns or concrete adjectives, its role in the representation of other semantic content (e.g. abstract non-event nouns such as “idea”) remains to be tested.

### Preferential encoding of object nouns in LIP, LPC and LIT

In a subset of cortical areas, including the LIP, LPC and LIT, we find that patterns of activity distinguished preferentially among concrete entity nouns than among verbs. These cortical areas are similar to those identified by previous work as being sensitive to distinctions among concrete noun. For example, Kumar and colleagues (2017) showed that patterns of activity in IP distinguish among different types of places (beaches, cities, highways and mountains). Analogously, Fairhall and Caramazza (2013) found that a portion of IT, extending into MTG, and the PC are sensitive to differences among mammals, birds, fruits, tools and clothes. The current results demonstrate that these regions are preferentially involved in representing the meanings of concrete nouns, as opposed to verbs.

Interestingly, prior research suggests that activity in these noun-responsive areas is sensitive to object category not only when participants are presented with words, but also when they are presented with pictures of objects (Simanova et al. 2012; Fairhall and Caramazza 2013; Devereux et al. 2013; Kumar et al. 2017). Furthermore, classifiers trained on patterns of activity produced by object nouns successfully decode among images of the same objects, and *vice versa* (Simanova et al. 2012; Fairhall and Caramazza 2013; Devereux et al. 2013; Kumar et al. 2017). One proposal is that these regions represent amodal conceptual knowledge that is not language-specific and can be accessed either through words or images (Binder and Desai 2011; Fairhall and Caramazza 2013). Notably, it remains to be tested whether these regions would also be engaged by images for which there are no lexical labels in the speaker’s language.

A key open question concerns the semantic dimensions that are represented within these concrete entity-responsive conceptual areas. Some clues come from the specific category distinctions to which these areas were sensitive in the current study. Within the noun category, we successfully decoded animals from places, and manmade from natural places. By contrast, the seemingly salient distinction between mammals and birds did not lead to distinguishable patterns of activity in these regions. Similarly, previous studies have found decoding between animals and inanimate objects (e.g. tools, places), as well as between manmade versus natural places (Simanova et al. 2012; Akama et al. 2012; Fairhall and Caramazza 2013; Devereux et al. 2013; Correia et al. 2014; Kumar et al. 2017). We are, however, not aware of any studies showing classification among animal nouns in these cortical areas. Why might this concrete entity-responsive cortical network be more sensitive to some semantic distinctions than others, particularly since participants easily make all the distinctions?

One hypothesis is that the spatial neural code within the concrete entity-responsive network more robustly reflects semantic distinctions that are highly inferentially relevant, such as animacy and the distinction between artifacts and natural kinds (Gelman 1983, 1988; Gelman and Markman 1986, 1987; Mandler et al. 1991; Bloom 1996; Gelman and Bloom 2000). Children distinguish animate and inanimate entities early in development (Gelman 1983; Mandler et al. 1991), and many languages mark animacy grammatically, distinguishing between entities that can and cannot serve as agents (Frawley 1992; Medin et al. 2000; Kemmerer 2017). Only animate entities engage in volitional behavior and have mental states, whereas inanimate entities are characterized by their functions, shape and compositional material (Frawley 1992). Analogously, the distinction of artifacts and natural kinds is conceptually significant, since only artifacts have intended functions and specific social roles (Bloom 1996; Diesendruck et al. 2003; Bromhead 2017).

Independent evidence for the idea that animacy is an organizing dimension of neural representations comes from neuropsychological studies showing that living and nonliving things dissociate in the context of brain damage (see Gainotti 2000; Capitani et al. 2003). Furthermore, animacy is a dimension that predicts localization at a coarse scale within the ventral visual stream during object recognition (Chao et al. 1999; Martin 2007; Krigeskorte et al. 2008; Mahon et al. 2009; Kanwisher 2010; Konkle and Caramazza 2013; Sha et al. 2015). An intriguing hypothesis is that the animacy dimension, which organizes the coarse spatial scale in the ventral visual stream, is represented at a finer spatial scale within the entity-responsive conceptual network. The development of spatial segregation along the dimension of animacy within these different cortical systems could be mutually constrained (Mahon and Caramazza 2011).

By contrast to animacy and the artifact/natural kind distinction, the distinction between birds and mammals is arguably less conceptually fundamental though equally salient. Children and adults often reason about living things based on knowledge of broad biological properties that are common to the animal class, such as breathes, eats, is born, sleeps and dies (Carey 1985). It has also been suggested that Western adults living in urban settings have sparse abstract knowledge about specific animal kinds (Medin and Atran 1999; Atran and Medin 2008; Medin and Bang 2014). When making fine-grained distinctions among animals of the kind participants were asked to make in the current study (e.g. elk vs. rhino), people often rely on physical appearance (e.g. shape, color) (Farah and McClelland 1991; Farah 1996). Such appearance related information may be stored outside of the concrete entity-responsive network identified in the current experiment. Previous studies suggest that parts of the ventral and dorsal object recognition streams are involved in representing appearance related information, including information that distinguishes between different animal types (Thompson-Schill et al. 1999; Oliver and Thompson-Schill 2003; Connolly et al. 2012, 2016). One possibility is that these representations were relied upon by the current participants only for the animal trials. Future studies could localize visual regions involved in animal recognition and ask whether these same regions are also involved in making similarity judgments about animals when using verbal stimuli.

In sum, one way to interpret the current findings is that not all perceptually salient distinctions between entities are equally likely to lead to detectable maps within conceptual neural networks. The abstract conceptual system may develop more robust maps of more central conceptual distinctions, such as agent *versus* inanimate entity and natural kind *versus* artifact. By contrast, distinctions among even the large animal classes, such as mammals and birds, may not be coded at a coarse enough spatial scale to be detectable with current MVPA techniques.

This null result should be interpreted with caution, however, since it is possible that differences among animal types are mapped in these cortical areas but were not detected in the current study, due to the nature of the task or other design elements resulting in insufficient power. In future work, it would be of interest to ask whether requiring participants to attend to abstract conceptual properties of mammals and birds (e.g. gives birth to live young vs. lay eggs) or preventing them from using perceptual information as a basis of judgment would improve decoding within the concrete entity-responsive network. It would also be interesting to ask whether animal experts (e.g. zoologists) would show more distinguishable patterns of activation for different animal types in these cortical areas.

Finally, it still remains possible that the putative abstract conceptual network actually distinguishes entities along perceptual dimensions (e.g. size, shape, color, texture), since all of the decodable categories do differ along such dimensions (e.g. places are much larger than animals). Future work should directly test the hypothesis that animacy and artifacts *versus* natural kinds are indeed dimensions that are encoded in this network, once perceptual characteristics are held constant.

### Distributed representation of verbs and nouns within the lexical-semantic network

While we find that verb- and noun-responsive regions have a bias toward representing semantic information related to their preferred word class, decoding was also successful for the non-preferred category in every cortical area tested. Analogous sensitivity to the non-preferred stimulus class has been observed in the visual object recognition literature. Images of objects from different classes (i.e. places, faces and bodies) can be distinguished from each other based on patterns of activity outside the traditional areas that preferentially respond to those categories (Haxby et al. 2001; Spiridon and Kanwisher 2002; Kanwisher and Yovel 2006). What does this sensitivity to the non-preferred lexical class reflect?

One possibility is that noun-related information is automatically retrieved during verb processing and *vice versa*. For example, retrieving a verb like “lick” may partially activate likely agents (e.g. dog) and objects (e.g. bone). A related possibility is that verb- and noun-responsive areas are involved in representing a type of semantic information that is particularly relevant for one grammatical class but is also relevant, to some degree, for the non-preferred category. For example, animacy is relevant to entities but also to agent of actions.

A further open question is how and whether activity in non-preferred cortical areas is functionally relevant to behavior. As noted in the introduction, neuropsychological evidence shows that verbs and nouns can dissociate in the context of brain damage (e.g. Goodglass et al. 1966; Luria and Tsvetkova 1967). This observation suggests that some neural populations are more behaviorally relevant for one grammatical class over another. Supporting evidence for this hypothesis also comes from studies with transcranial magnetic stimulation (TMS) (Papeo et al. 2014). Disruption of activity in the verb-responsive posterior LMTG region with TMS interferes with participants’ performance on a synonym-judgment task with verbs, but not nouns (Papeo et al. 2014). This observation suggests that despite being sensitive to distinctions among verbs and among nouns, verb-responsive LMTG is more behaviorally relevant for verb comprehension.

## Conclusions

We observed a double dissociation among cortical areas in their sensitivity to verb and noun semantics. An LMTG region that responds more to verbs than nouns is more sensitive to semantic distinctions among verbs than among nouns. By contrast, several parietal and inferior temporal areas (LPC, LIP and LIT) are more active during noun processing and more sensitive to distinctions among nouns. We hypothesize that the LMTG is preferentially involved in representing the meanings of verbs and events, while noun-responsive regions are preferentially involved in representing abstract conceptual information relevant to entities. However, all cortical areas tested were also sensitive to the semantic distinctions of their non-preferred grammatical class, albeit to a lesser degree. These results suggest that verb and noun meanings are represented in partially non-overlapping neural networks.

## Acknowledgments

We thank the F. M. Kirby Research Center for Functional Brain Imaging at the Kennedy Krieger Institute for their assistance with data collection; and the participants for making this research possible.

## References

Aggujaro S, Crepaldi D, Pistarini C, Taricco M, Luzzatti C. 2006. Neuro-anatomical correlates of impaired retrieval of verbs and nouns: Interaction of grammatical class, imageability and actionality. Journal of Neurolinguistics 19(3):175–94.

Akama H, Murphy B, Na L, Shimizu Y, Poesio M. 2012. Decoding semantics across fMRI sessions with different stimulus modalities: A practical MVPA study. Front Neuroinform 6: 24.

Anzellotti S and Caramazza A. 2017. Multimodal representations of person identity individuated with fMRI. Cortex 89: 85–97.

Atran S and Medin DL. 2008. The native mind and the cultural construction of nature. Cambridge (Mass.) [etc.: MIT Press.

Bedny M and Saxe R. 2012. Insights into the origins of knowledge from the cognitive neuroscience of blindness. Cognitive Neuropsychology 29(1–2):56–84.

Bedny M and Caramazza A. 2011. Perception, action, and word meanings in the human brain: The case from action verbs. Ann N Y Acad Sci 1224(1):81–95.

Bedny M and Thompson-Schill SL. 2006. Neuroanatomically separable effects of imageability and grammatical class during single-word comprehension. Brain Lang 98(2):127–39.

Bedny M, Dravida S, Saxe R. 2014. Shindigs, brunches, and rodeos: The neural basis of event words. Cognitive, Affective, & Behavioral Neuroscience 14(3):891–901.

Bedny M, McGill M, Thompson-Schill SL. 2008. Semantic adaptation and competition during word comprehension. Cerebral Cortex 18(11):2574–85.

Bedny M, Hulbert JC, Thompson-Schill SL. 2007. Understanding words in context: The role of Broca’s area in word comprehension. Brain Res 1146: 101–14.

Bedny M, Caramazza A, Pascual-Leone A, Saxe R. 2011. Typical neural representations of action verbs develop without vision. Cerebral Cortex 22(2):286–93.

Bedny M, Caramazza A, Grossman E, Pascual-Leone A, Saxe R. 2008. Concepts are more than percepts: The case of action verbs. J Neurosci 28(44):11347–53.

Berlin B, Breedlove DE, Raven PH. 1973. General principles of classification and nomenclature in folk biology. American Anthropologist 75(1):214–42.

Berlingeri M, Crepaldi D, Roberti R, Scialfa G, Luzzatti C, Paulesu E. 2008. Nouns and verbs in the brain: Grammatical class and task specific effects as revealed by fMRI. Cognitive Neuropsychology 25(4):528–58.

Bhat DNS. 2000. Word classes and sentential functions. Empirical Approaches to Language Typology:47–64.

Bi Y, Wang X, Caramazza A. 2016. Object domain and modality in the ventral visual pathway. Trends Cogn Sci (Regul Ed) 20(4):282–90.

Binder JR and Desai RH. 2011. The neurobiology of semantic memory. Trends Cogn Sci (Regul Ed) 15(11):527–36.

Bloom P. 1996. Intention, history, and artifact concepts. Cognition 60(1):1–29.

Bogka N, Masterson J, Druks J, Fragkioudaki M, Chatziprokopiou E, Economou K. 2003. Object and action picture naming in english and greek. European Journal of Cognitive Psychology 15(3):371–403.

Boylan C, Trueswell JC, Thompson-Schill SL. 2015. Compositionality and the angular gyrus: A multi-voxel similarity analysis of the semantic composition of nouns and verbs. Neuropsychologia 78: 130–41.

Bromhead H. 2017. The semantics of standing-water places in English, French, and Pitjantjatjara/Yankunytjatjara. In Ye, Z. (Ed.), The Semantics of Nouns. Oxford University Press.

Capitani E, Laiacona M, Mahon B, Caramazza A. 2003. What are the facts of semantic category-specific deficits? A critical review of the clinical evidence. Cognitive Neuropsychology 20(3–6):213–61.

Caramazza A and Hillis AE. 1991. Lexical organization of nouns and verbs in the brain. Nature 349(6312):788–90.

Carey S. 1985. Conceptual change in childhood. Cambridge, Mass.: MIT Press.

Carota F, Kriegeskorte N, Nili H, Pulvermüller F. 2017. Representational similarity mapping of distributional semantics in left inferior frontal, middle temporal, and motor cortex. Cerebral Cortex 27(1):294–309.

Chao LL, Haxby JV, Martin A. 1999. Attribute-based neural substrates in temporal cortex for perceiving and knowing about objects. Nat Neurosci 2(10).

Collina S, Marangolo P, Tabossi P. 2001. The role of argument structure in the production of nouns and verbs. Neuropsychologia 39(11):1125–37.

Connine CM, Mullennix J, Shernoff E, Yelen J. 1990. Word familiarity and frequency in visual and auditory word recognition. Journal of Experimental Psychology: Learning, Memory, and Cognition 16(6):1084.

Connolly AC, Guntupalli JS, Gors J, Hanke M, Halchenko YO, Wu YC, Abdi H, Haxby JV. 2012. The representation of biological classes in the human brain. J Neurosci 32(8):2608–18.

Connolly AC, Sha L, Guntupalli JS, Oosterhof N, Halchenko YO, Nastase SA, di Oleggio Castello MV, Abdi H, Jobst BC, Gobbini MI, et al. 2016. How the human brain represents perceived dangerousness or “predacity” of animals. J Neurosci 36(19):5373–84.

Cordier F, Croizet J, Rigalleau F. 2013. Comparing nouns and verbs in a lexical task. J Psycholinguist Res 42(1):21–35.

Correia J, Formisano E, Valente G, Hausfeld L, Jansma B, Bonte M. 2014. Brain-based translation: FMRI decoding of spoken words in bilinguals reveals language-independent semantic representations in anterior temporal lobe. J Neurosci 34(1):332–8.

Crepaldi D, Berlingeri M, Cattinelli I, Borghese NA, Luzzatti C, Paulesu E. 2013. Clustering the lexicon in the brain: A meta-analysis of the neurofunctional evidence on noun and verb processing. Front Hum Neurosci 7: 303.

Croft W. 2005. Word classes, parts of speech, and syntactic argumentation. Linguistic Typology 9(3):431–41.

Dale AM, Fischl B, Sereno MI. 1999. Cortical surface-based analysis: I. segmentation and surface reconstruction. Neuroimage 9(2):179–94.

Damasio AR and Tranel D. 1993. Nouns and verbs are retrieved with differently distributed neural systems. Proc Natl Acad Sci USA 90(11):4957–60.

Daniele A, Giustolisi L, Silveri MC, Colosimo C, Gainotti G. 1994. Evidence for a possible neuroanatomical basis for lexical processing of nouns and verbs. Neuropsychologia 32(11):1325–41.

Davis MH, Meunier F, Marslen-Wilson WD. 2004. Neural responses to morphological, syntactic, and semantic properties of single words: An fMRI study. Brain Lang 89(3):439–49.

de Diego Balaguer R, Rodríguez-Fornells A, Rotte M, Bahlmann J, Heinze H, Münte TF. 2006. Neural circuits subserving the retrieval of stems and grammatical features in regular and irregular verbs. Hum Brain Mapp 27(11):874–88.

den Ouden D, Fix S, Parrish TB, Thompson CK. 2009. Argument structure effects in action verb naming in static and dynamic conditions. Journal of Neurolinguistics 22(2):196–215.

Devereux BJ, Clarke A, Marouchos A, Tyler LK. 2013. Representational similarity analysis reveals commonalities and differences in the semantic processing of words and objects. Journal of Neuroscience 33(48):18906–16.

Diesendruck G, Hammer R, Catz O. 2003. Mapping the similarity space of children and adults’ artifact categories. Cognitive Development 18(2):217–31.

Downing PE, Jiang Y, Shuman M, Kanwisher N. 2001. A cortical area selective for visual processing of the human body. Science 293(5539):2470–3.

Earles JL and Kersten AW. 2000. Adult age differences in memory for verbs and nouns. Aging, Neuropsychology, and Cognition 7(2):130–9.

Engelkamp J, Zimmer H, Mohr G. 1990. Differential memory effects of concrete nouns and action verbs. Zeitschrift Für Psychologie Mit Zeitschrift Für Angewandte Psychologie.

Epstein R and Kanwisher N. 1998. A cortical representation of the local visual environment. Nature 392(6676):598–601.

Fairhall SL and Caramazza A. 2013. Brain regions that represent amodal conceptual knowledge. J Neurosci 33(25):10552–8.

Farah MJ. 1996. The living/nonliving dissociation is not an artifact: Giving an a priori implausible hypothesis a strong test. Cognitive Neuropsychology 13(1):137–54.

Farah MJ and McClelland JL. 1991. A computational model of semantic memory impairment: Modality specificity and emergent category specificity. J Exp Psychol: Gen 120(4):339.

Frawley W. 1992. Linguistic semantics. Hillsdale, NJ [u.a.]: Erlbaum.

Fujimaki N, Miyauchi S, Pütz B, Sasaki Y, Takino R, Tamada T. 1999. Functional magnetic resonance imaging of neural activity related to orthographic, phonological, and lexico - semantic judgments of visually presented characters and words. Hum Brain Mapp 8(1):44–59.

Gainotti G. 2000. What the locus of brain lesion tells us about the nature of the cognitive defect underlying category-specific disorders: A review. Cortex 36(4):539–59.

Garbin G, Collina S, Tabossi P. 2012. Argument structure and morphological factors in noun and verb processing: An fMRI study. PloS One 7(9):e45091.

Gelman R, Spelke ES, Meck E. 1983. What preschoolers know about animate and inanimate objects. In: The acquisition of symbolic skills. Springer. 297 p.

Gelman SA. 1988. The development of induction within natural kind and artifact categories. Cognit Psychol 20(1):65–95.

Gelman SA and Bloom P. 2000. Young children are sensitive to how an object was created when deciding what to name it. Cognition 76(2):91–103.

Gelman SA and Markman EM. 1987. Young children’s inductions from natural kinds: The role of categories and appearances. Child Dev:1532–41.

Gelman SA and Markman EM. 1986. Categories and induction in young children. Cognition 23(3):183–209.

Genovese CR, Lazar NA, Nichols T. 2002. Thresholding of statistical maps in functional neuroimaging using the false discovery rate. Neuroimage 15(4):870–8.

Gentner D. 1981. Some interesting differences between verbs and nouns. Cognition and Brain Theory 4: 161–78.

Glasser MF, Sotiropoulos SN, Wilson JA, Coalson TS, Fischl B, Andersson JL, Xu J, Jbabdi S, Webster M, Polimeni JR. 2013. The minimal preprocessing pipelines for the human connectome project. Neuroimage 80: 105–24.

Goodglass H, Klein B, Carey P, Jones K. 1966. Specific semantic word categories in aphasia. Cortex 2(1):74–89.

Green DM and Swets JA. 1966. Signal detection theory and psychophysics. New York: Wiley.

Grier JB. 1971. Nonparametric indexes for sensitivity and bias: Computing formulas. Psychol Bull 75(6):424.

Grindrod CM, Bilenko NY, Myers EB, Blumstein SE. 2008. The role of the left inferior frontal gyrus in implicit semantic competition and selection: An event-related fMRI study. Brain Res 1229: 167–78.

Grossman M, Koenig P, DeVita C, Glosser G, Alsop D, Detre J, Gee J. 2002. Neural representation of verb meaning: An fMRI study. Hum Brain Mapp 15(2):124–34.

Hanke M, Halchenko YO, Sederberg PB, Hanson SJ, Haxby JV, Pollmann S. 2009. PyMVPA: A python toolbox for multivariate pattern analysis of fMRI data. Neuroinformatics 7(1):37–53.

Haxby JV, Connolly AC, Guntupalli JS. 2014. Decoding neural representational spaces using multivariate pattern analysis. Annu Rev Neurosci 37: 435–56.

Haxby JV, Gobbini MI, Furey ML, Ishai A, Schouten JL, Pietrini P. 2001. Distributed and overlapping representations of faces and objects in ventral temporal cortex. Science 293(5539):2425–30.

Hernández M, Fairhall SL, Lenci A, Baroni M, Caramazza A. 2014. Predication drives verb cortical signatures. J Cogn Neurosci 26(8):1829–39.

Hillis AE and Caramazza A. 1995. Representation of grammatical categories of words in the brain. J Cogn Neurosci 7(3):396–407.

Jackendoff R. 1983. Semantics and cognition. MIT Press paperback ed. Cambridge, Mass.: MIT Press.

Jenkinson M, Bannister P, Brady M, Smith S. 2002. Improved optimization for the robust and accurate linear registration and motion correction of brain images. Neuroimage 17(2):825–41.

Kable JW, Lease-Spellmeyer J, Chatterjee A. 2002. Neural substrates of action event knowledge. J Cogn Neurosci 14(5):795–805.

Kable JW, Kan IP, Wilson A, Thompson-Schill SL, Chatterjee A. 2005. Conceptual representations of action in the lateral temporal cortex. J Cogn Neurosci 17(12):1855–70.

Kanwisher N. 2010. Functional specificity in the human brain: A window into the functional architecture of the mind. Proc Natl Acad Sci USA 107(25):11163–70.

Kanwisher N and Yovel G. 2006. The fusiform face area: A cortical region specialized for the perception of faces. Philos Trans R Soc Lond B Biol Sci 361(1476):2109–28.

Kanwisher N, McDermott J, Chun MM. 1997. The fusiform face area: A module in human extrastriate cortex specialized for face perception. J Neurosci 17(11):4302–11.

Kemmerer D. 2017. Categories of object concepts across languages and brains: The relevance of nominal classification systems to cognitive neuroscience. Language, Cognition and Neuroscience 32(4):401–24.

Kemmerer D, Castillo JG, Talavage T, Patterson S, Wiley C. 2008. Neuroanatomical distribution of five semantic components of verbs: Evidence from fMRI. Brain Lang 107(1):16–43.

Konkle T and Caramazza A. 2013. Tripartite organization of the ventral stream by animacy and object size. J Neurosci 33(25):10235–42.

Kriegeskorte N, Mur M, Ruff DA, Kiani R, Bodurka J, Esteky H, Tanaka K, Bandettini PA. 2008. Matching categorical object representations in inferior temporal cortex of man and monkey. Neuron 60(6):1126–41.

Kumar M, Federmeier KD, Fei-Fei L, Beck DM. 2017. Evidence for similar patterns of neural activity elicited by picture-and word-based representations of natural scenes. Neuroimage.

Laiacona M and Caramazza A. 2004. The noun/verb dissociation in language production: Varieties of causes. Cognitive Neuropsychology 21(2–4):103–23.

Langacker RW. 2008. Cognitive grammar: A basic introduction. Oxford University Press.

Langacker RW. 1987. Nouns and verbs. Language:53–94.

Lapinskaya N, Uzomah U, Bedny M, Lau E. 2016. Electrophysiological signatures of event words: Dissociating syntactic and semantic category effects in lexical processing. Neuropsychologia 93: 151–7.

Levin, B., Hovav, M.R.,. 1995. Unaccusativity at the syntax-lexical semantics interface. Linguistic Inquiry Monographs 26:ALL.

Levin, Beth,. 2007. English verb classes and alternations: A preliminary investigation. Chicago, Ill. [u.a.]: Univ. of Chicago Press.

Li P, Jin Z, Tan LH. 2004. Neural representations of nouns and verbs in Chinese: An fMRI study. Neuroimage 21(4):1533–41.

Liljeström M, Tarkiainen A, Parviainen T, Kujala J, Numminen J, Hiltunen J, Laine M, Salmelin R. 2008. Perceiving and naming actions and objects. Neuroimage 41(3):1132–41.

Luria AR and Tsvetkova LS. 1967. Towards the mechanisms of “dynamic aphasia”. Acta Neurol Psychiatr Belg 67(11):1045–57.

Luzzatti C, Raggi R, Zonca G, Pistarini C, Contardi A, Pinna G. 2002. Verb-noun double dissociation in aphasic lexical impairments: The role of word frequency and imageability. Brain Lang 81(1):432–44.

Mahon BZ and Caramazza A. 2011. What drives the organization of object knowledge in the brain? Trends Cogn Sci (Regul Ed) 15(3):97–103.

Mahon BZ, Anzellotti S, Schwarzbach J, Zampini M, Caramazza A. 2009. Category-specific organization in the human brain does not require visual experience. Neuron 63(3):397–405.

Mandler JM, Bauer PJ, McDonough L. 1991. Separating the sheep from the goats: Differentiating global categories. Cognit Psychol 23(2):263–98.

Marangolo P, Piras F, Galati G, Burani C. 2006. Functional anatomy of derivational morphology. Cortex 42(8):1093–106.

Martin A. 2007. The representation of object concepts in the brain. Annu Rev Psychol 58: 25–45.

Martin A, Haxby JV, Lalonde FM, Wiggs CL, Ungerleider LG. 1995. Discrete cortical regions associated with knowledge of color and knowledge of action. Science 270(5233):102.

Mätzig S, Druks J, Masterson J, Vigliocco G. 2009. Noun and verb differences in picture naming: Past studies and new evidence. Cortex 45(6):738–58.

McCarthy R and Warrington EK. 1985. Category specificity in an agrammatic patient: The relative impairment of verb retrieval and comprehension. Neuropsychologia 23(6):709–27.

Medin DL, Lynch EB, Solomon KO. 2000. Are there kinds of concepts? Annu Rev Psychol 51(1):121–47.

Medin DL and Scott A. 1999. Folkbiology. Cambridge (Mass.); London: MIT.

Medin DL and Bang M. 2014. Who’s asking?: Native science, western science, and science education. MIT Press.

Miceli G, Silveri MC, Nocentini U, Caramazza A. 1988. Patterns of dissociation in comprehension and production of nouns and verbs. Aphasiology 2(3–4):351–8.

Miceli G, Silveri MC, Villa G, Caramazza A. 1984. On the basis for the agrammatic’s difficulty in producing main verbs. Cortex 20(2):207–20.

Oliver RT and Thompson-Schill SL. 2003. Dorsal stream activation during retrieval of object size and shape. Cognitive, Affective, & Behavioral Neuroscience 3(4):309–22.

Palti D, Ben Shachar M, Hendler T, Hadar U. 2007. Neural correlates of semantic and morphological processing of Hebrew nouns and verbs. Hum Brain Mapp 28(4):303–14.

Papeo L, Lingnau A, Agosta S, Pascual-Leone A, Battelli L, Caramazza A. 2014. The origin of word-related motor activity. Cerebral Cortex 25(6):1668–75.

Pelphrey KA, Morris JP, Mccarthy G. 2004. Grasping the intentions of others: The perceived intentionality of an action influences activity in the superior temporal sulcus during social perception. J Cogn Neurosci 16(10):1706–16.

Pelphrey KA, Morris JP, Michelich CR, Allison T, McCarthy G. 2005. Functional anatomy of biological motion perception in posterior temporal cortex: An fMRI study of eye, mouth and hand movements. Cerebral Cortex 15(12):1866–76.

Perani D, Cappa SF, Schnur T, Tettamanti M, Collina S, Rosa MM, Fazio1 F. 1999. The neural correlates of verb and noun processing: A PET study. Brain 122(12):2337–44.

Peterson J. 2003. Languages without nouns and verbs? an alternative to lexical classes in Kharia. Old and new perspectives on south asian languages. grammar and semantics. papers growing out of the fifth international conference on south asian linguistics (ICOSAL-5), held at Moscow, Russia in July. 274 p.

Pollack I and Norman DA. 1964. A non-parametric analysis of recognition experiments. Psychonomic Science 1(1–12):125–6.

Rapp B and Caramazza A. 2002. Selective difficulties with spoken nouns and written verbs: A single case study. Journal of Neurolinguistics 15(3):373–402.

Rodd JM, Davis MH, Johnsrude IS. 2005. The neural mechanisms of speech comprehension: FMRI studies of semantic ambiguity. Cerebral Cortex 15(8):1261–9.

Sahin NT, Pinker S, Halgren E. 2006. Abstract grammatical processing of nouns and verbs in Broca’s area: Evidence from fMRI. Cortex 42(4):540–62.

Sapir E. 2004. Language: An introduction to the study of speech. Courier Corporation.

Sasse H. 1993. Syntactic categories and subcategories. Syntax: An International Handbook of Contemporary Research 2: 646–86.

Schreiber K and Krekelberg B. 2013. The statistical analysis of multi-voxel patterns in functional imaging. PLoS One 8(7):e69328.

Sha L, Haxby JV, Abdi H, Guntupalli JS, Oosterhof NN, Halchenko YO, Connolly AC. 2015. The animacy continuum in the human ventral vision pathway. J Cogn Neurosci.

Shapiro K and Caramazza A. 2003. Grammatical processing of nouns and verbs in left frontal cortex? Neuropsychologia 41(9):1189–98.

Shapiro K and Caramazza A. 2003. Looming a loom: Evidence for independent access to grammatical and phonological properties in verb retrieval. Journal of Neurolinguistics 16(2):85–111.

Shapiro K, Shelton J, Caramazza A. 2000. Grammatical class in lexical production and morhpological processing: Evidence from a case of fluent aphasia. Cognitive Neuropsychology 17(8):665–82.

Shapiro KA, Mottaghy FM, Schiller NO, Poeppel TD, Flüß MO, Müller H, Caramazza A, Krause BJ. 2005. Dissociating neural correlates for nouns and verbs. Neuroimage 24(4):1058–67.

Shapiro KA, Moo LR, Caramazza A. 2006. Cortical signatures of noun and verb production. Proc Natl Acad Sci USA 103(5):1644–9.

Simanova I, Hagoort P, Oostenveld R, Van Gerven MA. 2012. Modality-independent decoding of semantic information from the human brain. Cerebral Cortex 24(2):426–34.

Siri S, Tettamanti M, Cappa SF, Rosa PD, Saccuman C, Scifo P, Vigliocco G. 2007. The neural substrate of naming events: Effects of processing demands but not of grammatical class. Cerebral Cortex 18(1):171–7.

Smith SM, Jenkinson M, Woolrich MW, Beckmann CF, Behrens TE, Johansen-Berg H, Bannister PR, De Luca M, Drobnjak I, Flitney DE. 2004. Advances in functional and structural MR image analysis and implementation as FSL. Neuroimage 23:S208–19.

Spiridon M and Kanwisher N. 2002. How distributed is visual category information in human occipito-temporal cortex? an fMRI study. Neuron 35(6):1157–65.

Stanislaw H and Todorov N. 1999. Calculation of signal detection theory measures. Behavior Research Methods, Instruments, & Computers 31(1):137–49.

Stelzer J, Chen Y, Turner R. 2013. Statistical inference and multiple testing correction in classification-based multi-voxel pattern analysis (MVPA): Random permutations and cluster size control. Neuroimage 65: 69–82.

Swets JA, Tanner Jr WP, Birdsall TG. 1961. Decision processes in perception. Psychol Rev 68(5):301.

Szekely A, D’Amico S, Devescovi A, Federmeier K, Herron D, Iyer G, Jacobsen T, Anal’a LA, Vargha A, Bates E. 2005. Timed action and object naming. Cortex 41(1):7–25.

Tabossi P, Collina S, Caporali A, Pizzioli F, Basso A. 2010. Speaking of events: The case of CM. Cognitive Neuropsychology 27(2):152–80.

Thompson CK, Bonakdarpour B, Fix SC, Blumenfeld HK, Parrish TB, Gitelman DR, Mesulam M. 2007. Neural correlates of verb argument structure processing. J Cogn Neurosci 19(11):1753–67.

Thompson-Schill SL, Bedny M, Goldberg RF. 2005. The frontal lobes and the regulation of mental activity. Curr Opin Neurobiol 15(2):219–24.

Thompson-Schill SL, D’Esposito M, Kan IP. 1999. Effects of repetition and competition on activity in left prefrontal cortex during word generation. Neuron 23(3):513–22.

Thompson-Schill SL, D’Esposito M, Aguirre GK, Farah MJ. 1997. Role of left inferior prefrontal cortex in retrieval of semantic knowledge: A reevaluation. Proc Natl Acad Sci USA 94(26):14792–7.

Thompson-Schill SL, Swick D, Farah MJ, D’Esposito M, Kan IP, Knight RT. 1998. Verb generation in patients with focal frontal lesions: A neuropsychological test of neuroimaging findings. Proc Natl Acad Sci USA 95(26):15855–60.

Tranel D, Grabowski TJ, Lyon J, Damasio H. 2005. Naming the same entities from visual or from auditory stimulation engages similar regions of left inferotemporal cortices. J Cogn Neurosci 17(8):1293–305.

Tsao DY and Livingstone MS. 2008. Mechanisms of face perception. Annu Rev Neurosci 31: 411–37.

Tyler LK, Stamatakis E, Dick E, Bright P, Fletcher P, Moss H. 2003. Objects and their actions: Evidence for a neurally distributed semantic system. Neuroimage 18(2):542–57.

Tyler LK, Randall B, Stamatakis EA. 2008. Cortical differentiation for nouns and verbs depends on grammatical markers. J Cogn Neurosci 20(8):1381–9.

Tyler LK, Bright P, Fletcher P, Stamatakis EA. 2004. Neural processing of nouns and verbs: The role of inflectional morphology. Neuropsychologia 42(4):512–23.

Tyler LK, Russell R, Fadili J, Moss HE. 2001. The neural representation of nouns and verbs: PET studies. Brain 124(8):1619–34.

Vander Wyk BC, Hudac CM, Carter EJ, Sobel DM, Pelphrey KA. 2009. Action understanding in the superior temporal sulcus region. Psychological Science 20(6):771–7.

Vigliocco G, Vinson DP, Druks J, Barber H, Cappa SF. 2011. Nouns and verbs in the brain: A review of behavioural, electrophysiological, neuropsychological and imaging studies. Neuroscience & Biobehavioral Reviews 35(3):407–26.

Walther DB, Caddigan E, Fei-Fei L, Beck DM. 2009. Natural scene categories revealed in distributed patterns of activity in the human brain. J Neurosci 29(34):10573–81.

Walther DB, Chai B, Caddigan E, Beck DM, Fei-Fei L. 2011. Simple line drawings suffice for functional MRI decoding of natural scene categories. Proc Natl Acad Sci USA 108(23):9661–6.

Wang J, Baucom LB, Shinkareva SV. 2013. Decoding abstract and concrete concept representations based on single - trial fMRI data. Hum Brain Mapp 34(5):1133–47.

Watson DM, Hartley T, Andrews TJ. 2014. Patterns of response to visual scenes are linked to the low-level properties of the image. Neuroimage 99: 402–10.

Watson DM, Hartley T, Andrews TJ. 2017. Patterns of response to scrambled scenes reveal the importance of visual properties in the organization of scene-selective cortex. Cortex 92: 162–74.

Weide RL. 1998. The CMU pronouncing dictionary. URL: Http://www.Speech.Cs.Cmu.edu/cgibin/cmudict.

Yu X, Bi Y, Han Z, Zhu C, Law S. 2012. Neural correlates of comprehension and production of nouns and verbs in chinese. Brain Lang 122(2):126–31.

Yu X, Law SP, Han Z, Zhu C, Bi Y. 2011. Dissociative neural correlates of semantic processing of nouns and verbs in Chinese—A language with minimal inflectional morphology. Neuroimage 58(3):912–22.

Zingeser LB and Berndt RS. 1990. Retrieval of nouns and verbs in agrammatism and anomia. Brain Lang 39(1):14–32.

